# The mRNA export adaptor Yra1 contributes to DNA double-strand break repair through its C-box domain

**DOI:** 10.1101/441980

**Authors:** Valentina Infantino, Evelina Tutucci, Noël Yeh Martin, Audrey Zihlmann, Varinia García-Molinero, Géraldine Silvano, Benoit Palancade, Françoise Stutz

## Abstract

Yra1 is an mRNA export adaptor involved in mRNA biogenesis and export in *S. cerevisiae*. Yra1 overexpression was recently shown to promote accumulation of DNA:RNA hybrids favoring DNA double strand breaks (DSB), cell senescence and telomere shortening, via an unknown mechanism. Yra1 was also identified at an HO-induced DSB and Yra1 depletion causes defects in DSB repair. Previous work from our laboratory showed that Yra1 ubiquitination by Tom1 is important for mRNA export. Interestingly, we found that Yra1 is also ubiquitinated by the SUMO-targeted ubiquitin ligases Slx5-Slx8 implicated in the interaction of irreparable DSB with nuclear pores. Here we show that Yra1 binds an HO-induced irreparable DSB. Importantly, a Yra1 mutant lacking the evolutionarily conserved C-box is not recruited to an HO-induced irreparable DSB and becomes lethal under DSB induction in a HO-cut reparable system. Together, the data provide evidence that Yra1 plays a crucial role in DSB repair via homologous recombination. Unexpectedly, while the Yra1 C-box is essential, Yra1 sumoylation and/or ubiquitination are dispensable in this process.

## INTRODUCTION

Yra1 (<u>Y</u>east <u>R</u>NA <u>a</u>nnealing protein <u>1</u>) is an essential protein in *S. cerevisiae*, well characterized as an mRNA export adaptor involved in transcription elongation, 3’ processing, and finally mRNA export together with the Mex67/Mtr2 export receptor and the poly(A) binding protein Nab2 (1).

Yra1 is evolutionarily conserved from yeast to human and belongs to the <u>R</u>NA and <u>E</u>xport <u>F</u>actor (REF) family of hnRNP-like proteins (2-5). REF proteins include a conserved domain organization with a central RNP-motif containing an RNA binding domain (RBD) and two highly conserved N-and C-terminal boxes (N-box and C-box). These domains are separated by two variable regions (N-var and C-var), rich in positively charged amino acids that mediate interaction with RNAs and Mex67 (3, 6). *yra1* mutants lacking the RBD, the N-terminal or C-terminal (N-box+N-var or C-box+C-var) regions are viable indicating functional redundancy in their RNA binding properties. However, at least one highly conserved N-box or C-box is required for viability as deletion of both is lethal (7).

While Mex67 and Nab2 are shuttling between the nucleus and cytoplasm, Yra1 is a strictly nuclear protein (3, 5). Nuclear localization of Yra1 is important for mRNA export as mutants lacking the N-terminal nuclear localization signal (NLS) demonstrate nuclear accumulation of poly(A)+ RNA when examined by fluorescent *in situ* hybridization (FISH) (7). Yra1 binding to mRNA is important as well, as *yra1* mutants lacking the N-or C-terminal variable RNA binding domains also present poly(A)+ RNA export defects.

Interestingly, *yra1* mutants lacking the RBD also show some poly(A)+ RNA export defect although this domain is not implicated in Mex67 interaction nor RNA binding *in vitro*, suggesting that it may contribute to Yra1 function by ensuring optimal folding of the protein. Loss of the highly conserved Yra1 C-box (*yra1(1-210)* mutant) does not cause an obvious poly(A)+ mRNA export defect, but it is required for optimal growth (7). This observation is consistent with the fact that the C-box does not play a major role in Mex67 or RNA binding and suggests that this highly conserved 16 amino acids sequence may be important for another aspect of Yra1 function.

Different layers of regulations have been shown to modulate Yra1 levels and function in mRNA biogenesis. We have previously shown that Yra1 ubiquitination by the E3 ligase Tom1 displaces Yra1 from messenger ribonucleoparticles (mRNPs) as a quality control signal for correctly processed mRNP prior to export into the cytoplasm (8). Another important feature for Yra1 regulation is that the *YRA1* gene harbors the second largest intron (776 nt) in the *S. cerevisiae* genome, containing a non-canonical <u>b</u>ranchpoint <u>s</u>equence (BS, gACUAAC) after a long first exon (300 nt). An excess of Yra1 protein prevents *YRA1* pre-mRNA splicing and promotes export of the unspliced transcript into the cytoplasm where it is degraded by the 5’ to 3’ decay pathway (4, 9). It has been reported previously that the *yra1∆intron* mutant shows Yra1 protein overexpression (7) that is toxic for cell growth (10, 11) and impairs poly(A)+ RNA export (12, 13). The presence of the *YRA1* intron is important to maintain optimal Yra1 protein levels through Yra1 auto-regulation at the level of splicing in a negative feedback mechanism (4). Studies on the *YRA1* gene revealed that a mutation restoring a canonical branch-point sequence in the context of the C-terminal mutation *yra1-F223S* or the C-terminal deletion *yra1-ΔC11* is not viable. Overall, these mutagenesis experiments indicated that at least three elements contribute to optimal Yra1 autoregulation: a long first exon, a long intron, a weak branchpoint and an intact C-terminal domain (4). The C-terminal domain was proposed to negatively regulate splicing provided that splicing efficiency was suboptimal.

Independent studies have also indicated that besides its function in mRNA biogenesis and export, Yra1 could contribute to DNA metabolism. It was initially proposed that Yra1 interacts with a subunit of the DNA polymerase δ and Dia2, an E3 ubiquitin ligase involved in DNA replication, genome stability and S phase checkpoint recovery (14, 15). The C-box domain of Yra1 was suggested to be necessary for the recruitment of Dia2 at replication origins (15), establishing a potential functional link between Yra1 and DNA metabolism. Another report provided evidence that strong Yra1 overexpression causes transcription-associated hyper-recombination, a cell senescence-like phenotype and telomere shortening, probably by counteracting telomere replication since overexpressed Yra1 was located at the Y telomeric regions by ChIP-chip. In the proposed model, Yra1 overexpression stabilizes R-loops, which contain DNA:RNA hybrids and displaced DNA strands, favoring conflicts between the replication fork and RNA Pol II resulting in genome instability (16, 17). Finally, a recent study based on ChAP-MS (chromatin affinity purification with mass spectrometry) identified Yra1 in association with a reparable double-strand break (DSB); moreover, a *yra1* DAmP (Decreased Abundance by mRNA Perturbation) hypomorph mutant showed sensitivity to DSB agents and global defects in DSB repair by pulse field gel electrophoresis (18).

DSBs can be repaired by two independent pathways: non-homologous end joining (NHEJ) that joins the DNA ends of the lesion in an error-prone process, and homologous recombination (HR), an error-free pathway used when homologous DNA sequences are available for the repair (19). The genetic instability resulting from unrepaired DSBs leads to cell death (20). The HR process has to be tightly regulated to avoid aberrant genomic rearrangements. On site sumoylation of HR proteins induced under DNA damage is pivotal to ensure efficient and optimal DSB repair (21-24). SUMO-targeted E3 ubiquitin ligases (STUbL), such as the Slx5-Slx8 complex in yeast, have also been shown to contribute to the maintenance of genome stability, although their targets have not been systematically identified (25).

In this work, we show that Yra1 is sumoylated by the SUMO ligases Siz1 and Siz2, desumoylated by the SUMO protease Ulp1 and ubiquitinated by the SUMO-dependent E3 ligases Slx5-Slx8, which are important for genome integrity (25, 26). Importantly, we find that Yra1 is recruited to DSBs and identify the Yra1 C-box domain to be crucial for the binding and repair; however, Yra1 ubiquitination and/or sumoylation are not required in this process. Our results strengthen the importance of Yra1 in genome integrity and provide evidence for a critical role of Yra1 in DSB repair.

## MATERIALS AND METHODS

### Yeast strains and plasmids

The strains and plasmids used in this study are listed in Supplementary Tables 1 and 2 (S1 and S2 Table). Primers are listed in Supplementary Table 3 (S3 Table).

The *YRA1* shuffled strains were obtained by transformation of the *YRA1* shuffle strain (*yra1::HIS3*, YCpLac33-URA3-*YRA1*WT, Cen) with YCpLac22-TRP centromeric plasmids encoding wild-type HA-tagged Yra1. The transformed strains were plated on 5-FOA to select against the WT *YRA1* URA3 plasmid. The cells able to grow on 5-FOA contain only the YCplac22-TRP1*-*HA*-YRA1* WT plasmid (*YRA1* shuffled background). Single clones were analyzed for correct auxotrophic markers and checked for HA-Yra1 expression by Western blot with αHA antibodies.

The strains with integrated WT or mutant *HA-YRA1* were obtained by transformation of the W303 Mat-a/α diploid strain or FSY5073 (GA-6844 HO irreparable system (27)) with a fragment containing the HA-tagged wild-type or mutant *YRA1* sequences obtained by SmaI digestion of an engineered pUC18 construct. The pUC18 plasmids were obtained by Gibson assembly and contain a SmaI fragment consisting of the HA-tagged wild-type or mutant *YRA1* sequences preceded by the *YRA1* promoter and followed by the *YRA1* 3’ UTR, a selective marker (URA3 or HIS3) and an additional 100 pb of *YRA1* 3’ downstream sequences. Yeast transformants were plated on the relevant selective medium. Correct recombination and integration into the endogenous *YRA1* locus was checked by PCR with a forward primer complementary to a sequence −600bp upstream of the *YRA1* locus (OFS3118), not present in the plasmid sequence, and a reverse primer matching the HA-tag sequence present only in the plasmid-derived sequence (OFS3120). The W303 diploid strains containing the integrated *HA-YRA1 WT* or *HA-yra1* mutant sequences were sporulated on K-acetate agar plates for 3 days at 25°C and dissected. Single spores were analyzed for relevant auxotrophic markers; HA-*YRA1* integration was confirmed by PCR as described above and expression of HA-Yra1 proteins was verified by Western blot.

The deletion strains were generated by homologous recombination of a cassette containing an auxotrophic marker flanked by sequences adjacent to the gene to delete. The pUG73::*LEU2,* pAG25::natMX4 or pUG6::kanMX6 cassettes were amplified by PCR using 80 nucleotides long forward and reverse primers (20 nt complementary to the plasmid and 60 nt complementary to the target sequences). PCR products were transformed into the *YRA1* shuffle or WT W303 Mat-a/α diploid strains and correct insertion confirmed by PCR. The W303 Mat-a/α diploid strains containing the gene deletion were sporulated and single spores analyzed for auxotrophic markers. Haploid Mat-α WT W303 deletion mutants were crossed with haploid Mat-a strains containing integrated HA-YRA1 WT or mutant sequences obtained as described above. The diploid *yra1* double mutants were sporulated to obtain haploid *yra1* double mutants in W303 background. In the case of deletions in the *YRA1* shuffle, the *yra1* double mutants were obtained by plasmid shuffling as explained above.

The strains with integrated *HA-YRA1* in FSY6881 (NA17 strain with HO reparable system) (28) were obtained after four back crosses between the integrated *HA-YRA1 WT* or *HA-yra1(1-210)* and *HA-yra1allKR* mutants in W303 and the NA17 strain. The sporulation, dissection and analysis of the strains was performed as described above. The presence of the cassette KanMX::HO-cs at URA3 and KanMX::ClaI at LYS2 was checked by PCR followed by digestion with the restriction enzymes BamH1 (near the HO site) and ClaI.

### Media and culture conditions

If not specified, yeast strains were thawed on yeast extract-peptone-dextrose (YPD) plates and grown for two days at 25°C. Cells were pre-cultured in 5 ml of liquid YPD to reach an OD_600_= 0.7-0.8 at 25°C and diluted into 100 ml YPD overnight culture to reach OD_600_= 0.8-1 at 25°C in the morning.

For the protein stability assays using metabolic depletion of *GAL-HA-YRA1* in presence of the endogenous wild-type *YRA1* gene, cells expressing HA-Yra1 from the GAL promoter on a centromeric plasmid were grown over-night in selective medium containing 2% galactose. When reaching OD_600_=0.3, cells were shifted to selective medium containing 2% glucose to repress *GAL-HA-YRA1* and collected at time 0, 1h, 2h, 3h, 4h, 5h, 6h, and 7h following glucose addition.

To induce the HO endonuclease-mediated irreparable DSB, cells were grown over-night in SCLGg (SC lactate 2%/glycerol 2% containing 0.05% Glucose). Cells at OD_600_=0.4 were shifted to SCLGg medium containing 2% glucose for 2h (no cut induction) or to SCLGg medium containing 2% galactose to induce the HO endonuclease. Cells were collected at 30 minutes, 1h, 2h and 4h following galactose addition.

To induce the HO endonuclease-mediated reparable DSB, cells were grown over-night in SCLGg (SC lactate 2%/glycerol 2% containing 0.05% Glucose). Exponentially growing cells were treated with 2% galactose to induce the HO endonuclease or not (control) for 2h. Serial dilutions of 200/100/50 cells were plated on SCLGg Glu 2%. In another related experiment, serial dilutions of exponentially growing cells in SCLGg medium were directly plated on SCLGg Gal 2% or SCLGg Gal 3%-Raf 1% to induce the HO cut, and on SCLGg Glu 2% to repress HO endonuclease expression.

### Spot test

Cells grown in YPD medium to stationary phase were diluted to OD_600_=1 and five 10-fold serial dilutions were prepared for spotting on agar plates. For each spot, 3μl were deposited on 2% glucose YPD plates in the presence or absence of drug (Zeocin 25 μg/ml, 50 μg/ml, and 100 μg/ml). Plates were incubated at 25°C, 30°C, 34°C or 37°C for 3 days.

### Protein extraction and Western blotting

Cells were grown to OD_600_=1. Cell lysis was performed by adding 1 ml H_2_O with 150μl of Yex-lysis buffer (1.85M NaOH, 7.5% 2-mercaptoethanol) to the pellet of 5 ODs of cells and kept 10 minutes on ice. Proteins were precipitated by addition of 150μl of TCA 50% for 10 minutes on ice. The pellet was resuspended in 30μl of 1X sample buffer (1M Tris-HCl pH6.8, 8 M Urea, 20% SDS, 0.5M EDTA, 1% 2-mercaptoethanol, 0.05% bromophenol blue). Total protein extracts were fractioned on SDS-PAGE and examined by Western blotting with αHA (Enzo), αYra1 (Stutz laboratory), αPgk1 (Abcam), αRfa1, 2, 3 to detect RPA (kind gift from Vincent Géli), αGFP (Roche), αRad51 (Abcam) antibodies. For quantitative Western blot analyses, fluorescent secondary α-Mouse (IRDye 800CW) and α-Rabbit (IRDye 680RD) antibodies were used. The signals were revealed with the LI-COR instrument.

### Chromatin immunoprecipitation (ChIP) and quantitative real-time PCR

Cells grown to OD_600_=1 were cross-linked with 1.2% of formaldehyde (Molecular Biology grade Calbiochem^TM^) for 10 minutes at 25°C under continuous gentle agitation, quenched with 250mM of glycine (Sigma) for 5 min at 25°C and then on ice for at least 5 min, washed with PBS 1X and frozen at −20°C. Pellets of 100 ml cultures at OD_600_=1 were resuspended in 1ml of FA lysis buffer (10mM HEPES KOH pH 7.5, 140mM NaCl, 1mM EDTA pH 8, 1%Triton X-100, 0.1% sodium deoxycholate) containing a protease inhibitor cocktail (cOmplete tablets, Mini EDTA-free, Roche). Cells were mechanically broken with a magnalyser at 6500rpm for 30 seconds (4 times), and genomic DNA was sonicated for 20 cycles of 30 seconds ON/OFF in presence of 0.5% SDS added before the sonication step. Samples were centrifuged at 13000rpm for 15 min at 4°C, and chromatin (supernatant phase) was quantified by Bradford. For each IP, 1/10 of the total extract was kept as INPUT for final normalization. Chromatin extracts (500μg) were incubated at 4°C o/n with a specific antibody. In parallel, magnetic beads (Dynabeads® Magnetic, Thermo Fisher Scientific) were incubated with BSA 5 mg/ml at 4°C o/n. The magnetic beads were washed twice with FA lysis buffer and resuspended with the same volume of FA lysis buffer containing a protease inhibitor cocktail (beads 50% v/v). The chromatin extracts with a specific antibody were incubated with 30μl of magnetic beads for 4h at 4°C on a rotating wheel. The magnetic beads were then washed twice with FA lysis buffer, twice with FA 500 (50mM HEPES KOH pH 7.5, 500mM NaCl, 1mM EDTA pH 8, 1%Triton X-100, 0.1% sodium deoxycholate), once with Buffer III (20mM Tris-HCl pH 8, 1mM EDTA pH 8, 250mM LiCl, 0.5% NP40, 0.5% sodium deoxycholate) and once with TE 1X (100mM Tris-HCl pH 8, 10mM EDTA pH 8). DNA was eluted with 200μl of elution buffer (50mM Tris-HCl pH 7.5, 1% SDS) at 65°C for 20 minutes. IP and INPUT DNAs were finally de-crosslinked with proteinase K (Roche) (0.4 μg/μl) for 2 hours at 42°C, and o/n at 65°C. The decrosslinked IP and INPUT DNAs were purified (Promega, Wizard® Genomic DNA Purification Kit). IP and INPUT (2μl) were quantified by qPCR with SYBR® Green PCR Master Mix (Applied Biosystems) using specific primers.

The following antibodies were used: a rabbit polyclonal αHA antibody (Enzo), a rabbit polyclonal αYra1 antibody and corresponding pre-immune (Stutz laboratory).

### Ubiquitination and Sumoylation assays

Ubiquitination and sumoylation assays were performed essentially as described (8, 29, 30) using cells transformed with a plasmid expressing His6-Ubi or His6-SUMO from a copper inducible promoter (*P_CUP1_*). Briefly Ubiquitin/SUMO expression was induced with 0.1 mM CuSO_4_ overnight or for 3h. Cell cultures (200 ml) at OD_600_=1 were collected adding TCA 5% for 20 minutes to allow protein precipitation. Cell pellets were washed twice with acetone 100%. Dry pellets were resuspended with 1ml of Guanidinium buffer (100 mM sodium phosphate at pH 8, 10 mM Tris-HCl, 6 M guanidinium, 10 mM imidazole, 0.2% Triton X-100, 10 mM NEM, complete protease inhibitor mix [Roche]) prior to cell disruption with glass beads in a magnalyser (6 cycles at 6500 rpm for 1 minute).

Cells lysates were spun at 13000 rpm for 20 min. Between 5-8 mg of protein from the supernatant was incubated with 100μl of Ni-NTA acid-agarose (Qiagen) for 2h at room temperature on a rotating wheel. Agarose beads were washed once with Guanidinium buffer and three times with Urea buffer (100 mM sodium phosphate at pH 6.8, 10 mM Tris-HCl, 8M urea, 20 mM imidazole, 0.2% Triton X-100, complete protease inhibitor mix [Roche]). His6-ubiquitinated and His6-SUMOylated proteins were eluted with 40 μl of Sample Buffer and boiled for 5 min at 95°C. 20μl samples were analyzed by Western blot with the relevant antibodies: αHIS for ubiquitinated or SUMOylated proteins, αHA for ubiquitinated or SUMOylated HA-Yra1. Input samples were also precipitated with TCA 5%, the pellets resuspended with Sample Buffer and boiled 5 min at 95°C to be analyzed by Western Blot with αHA for HA-Yra1 and αPgk1 as loading control.

### FISH experiments

The FISH experiments on the *YRA1* shuffled strains deleted for various ubiquitin ligases were done essentially as described in (8), while the FISH experiments on the different HA-YRA1 integrated strains were performed as described in (31). In the latter case, images were taken with the Zeiss LSM700 confocal microscope using laserline 405 nm for DAPI detection and laserline 555 nm for Cy3. Transmission light images were taken to see the cell shape. In both cases Poly(A)^+^mRNA in situ hybridization was performed with a Cy3-labeled oligo-dT_(50)_ probe.

### Colony Forming Unit Assay (CFU)

Three serial dilutions (200/100/50) of exponentially growing cells in SCLGg medium were plated on SCLGg Gal 2% or SCLGg Gal 3%-Raf 1% or SCLGg Glu 2% and incubated at 25°C for 5 days. CFUs were counted and the % of colonies was expressed as CFU relative to the CFU grown on SCLGg Glu 2% (control).

## RESULTS

### Yra1 is modified by the SUMO-targeted E3 ubiquitin ligase Slx5-Slx8

Previous work from our laboratory showed that Yra1 ubiquitination by Tom1 elicits Yra1 dissociation from mRNPs, presumably in the context of the nuclear pore complex (NPC), allowing proper mRNP export into the cytoplasm (8). Intriguingly, Yra1 ubiquitination is not fully abrogated in the *Δtom1* mutant, suggesting that other E3 ligases are involved in Yra1 regulation, possibly for other Yra1 functions. In view of the putative role of Yra1 in genome stability, we wondered whether this protein could be modified by SUMO-dependent ubiquitination. Consistently, we identified the SUMO-targeted E3 ubiquitin ligase (STUbL) complex Slx5-Slx8 to be responsible for Yra1 ubiquitination together with Tom1 (**Fig 1**).

**Fig 1:**
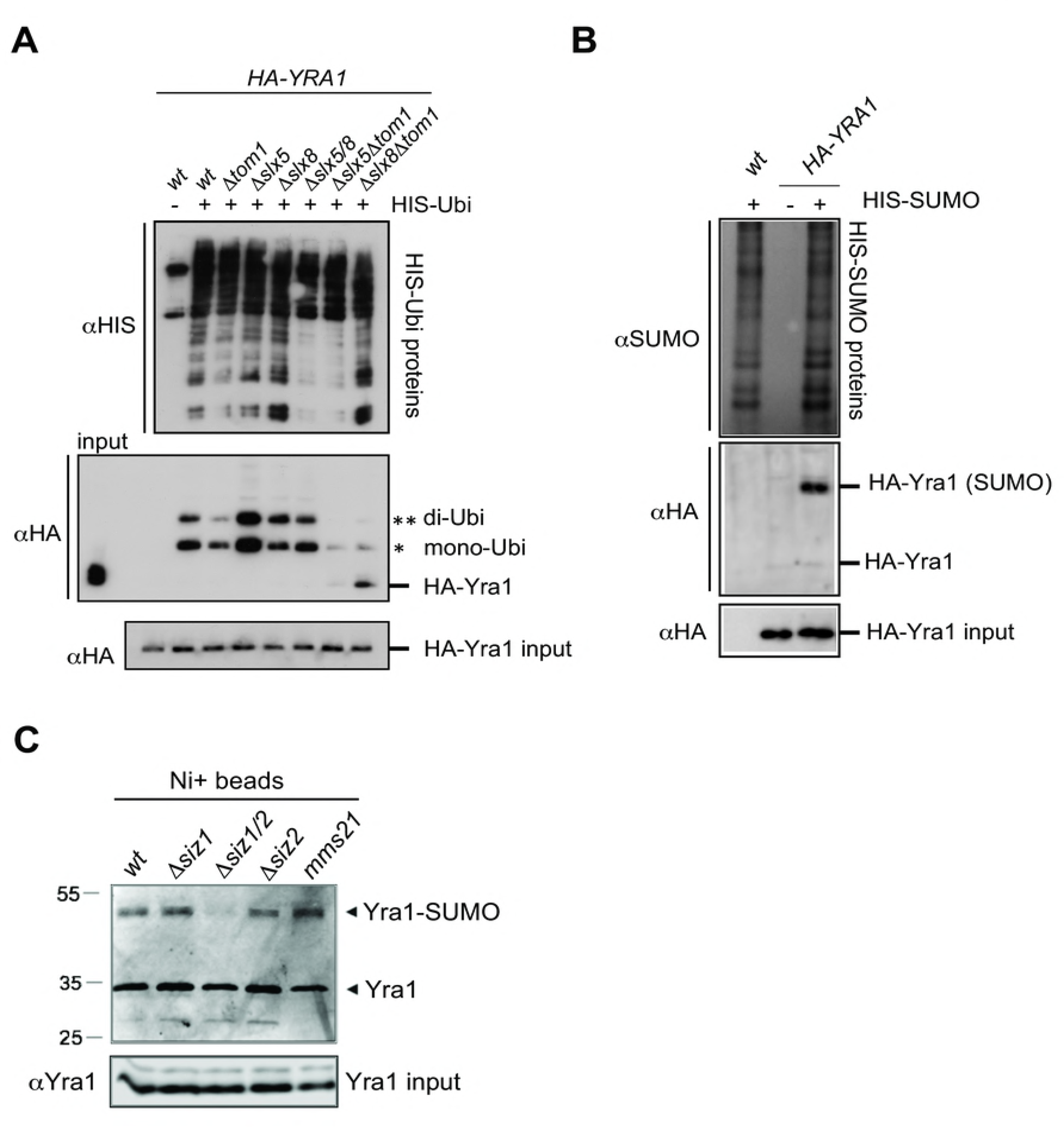
Yra1 is a sumoylated protein targeted for ubiquitination by the SUMO-dependent ubiquitin ligase Slx5-8. **(A)** Yra1 ubiquitination depends on the STUbL Slx5-Slx8 and Tom1. Ubiquitination assay of shuffled *HA-YRA1* in wild-type and in *Δtom1*, *Δslx8*, *Δslx5*, *Δslx8Δtom1*, *Δslx5Δtom1* and *Δslx8Δslx5* mutant backgrounds. Strains were transformed with a copper inducible His-Ubiquitin expressing 2μ plasmid (+) or an empty vector (-). His-Ubiquitin was induced with copper over night. His-Ubiquitinated proteins were affinity-purified and the Ubiquitinated forms of Yra1 detected by Western Blot with αHA antibodies. Western Blot of input samples with αHA antibody was used to assess input Yra1 levels. One representative experiment of 3 is shown. **(B)** Yra1 is sumoylated. Sumoylation assay in wild-type and *HA-YRA1* backgrounds. Strains were transformed with a copper inducible His-SUMO expressing 2μ plasmid (+) or with an empty vector (-). His-SUMO was induced with copper for 3h. His-sumoylated proteins were affinity-purified and the sumoylated forms of Yra1 detected by Western Blot with αHA antibodies. Western Blot of input samples with αHA was used to assess input Yra1 levels. One representative experiment of 3 is shown. **(C)** Yra1 is sumoylated by Siz1/Siz2. Yra1 sumoylation assay in wild-type as well as *Δsiz1*, *Δsiz1/siz2*, *Δsiz2* and *mms21* mutant backgrounds. Strains were transformed with a copper inducible His-SUMO expressing 2μ□plasmid. His-SUMO was induced with copper for 3h. His-SUMOylated proteins were affinity purified and the SUMOylated forms of Yra1 detected by Western Blot with an αYrα1 antibody. Western Blot of input samples with αYra1 was used to assess input Yra1 levels. One representative experiment of 2 is shown.

The ubiquitination assay of HA-Yra1 in wild-type and in *Δtom1*, *Δslx5*, *Δslx8*, *Δslx5Δslx8*, *Δslx5Δtom1*, *Δslx8Δtom1* mutant backgrounds showed that the Yra1 ubiquitination detected in the *Δtom1* mutant was completely abrogated in the *Δslx5Δtom1* and *Δslx8Δtom1* double mutants (**Fig 1A**), indicating a role for both the Slx5-Slx8 and Tom1 E3 ligases in Yra1 regulation. Since the Slx5-Slx8 E3 ligase complex is stimulated by substrate sumoylation (32), and in view of the reported identification of Yra1 as potentially sumoylated in a proteome-wide study (33), we confirmed that Yra1 is indeed sumoylated (**Fig 1B** and **1C**). Both Siz1 and Siz2 SUMO E3 ligases are involved in this modification as Yra1 sumoylation is fully abrogated in the *Δsiz1Δsiz2* double mutant background (**Fig 1C**). Furthermore, Yra1 is de-sumoylated by the SUMO protease Ulp1 as Yra1 sumoylation increased in the *ulp1* <u>t</u>emperature-<u>s</u>ensitive (*ts*) mutant (**Supplementary Fig S1A**). These data support the hypothesis that Yra1 is regulated both by sumoylation and ubiquitination. In addition, HA-Yra1 ubiquitination was increased in the *ulp1 ts* mutant compared to a wild-type background, suggesting a possible stimulating effect of sumoylation on ubiquitination (**Supplementary Fig S1B**).

Known targets of Slx5-Slx8 are controlled by ubiquitin-dependent proteasomal degradation (34-38). To define whether Yra1 ubiquitination by Slx5-Slx8 may target Yra1 to degradation, we used metabolic depletion to examine Yra1 turnover. Because *YRA1* is essential, an HA-tagged version of *YRA1* was expressed from a galactose-inducible promoter on a plasmid transformed into a strain expressing a wild-type *YRA1* gene. Switching cells from galactose to glucose-containing medium represses *GAL-HA-YRA1* gene expression and allows following the decay of the HA-Yra1 protein in different genetic backgrounds. Under metabolic glucose repression, HA-Yra1 has a half-life of 3.8h (**Supplementary Fig S2A**). No significant stabilization of HA-Yra1 protein was detected in *Δslx8, Δslx5, Δtom1* or *Δslx8Δtom1* (**Supplementary Fig S2B and S2C**), suggesting that ubiquitination by Slx5-Slx8 does not lead to Yra1 degradation by the proteasome.

We previously proposed that Yra1 regulation by Tom1 is linked to the function of Yra1 in mRNP export (1, 8). Visualization of poly(A)+ RNA distribution by fluorescence in situ hybridization (FISH) in the *Δslx5* and *Δslx8* single mutants did not show any nuclear poly(A)+ RNA retention while the *Δslx5Δtom1* (32.3%) and *Δslx8Δtom1* (26%) double mutants had mRNA export defects comparable to the *Δtom1* mutant (30.8%) (**Supplementary Fig S3A**). These observations suggest that Yra1 ubiquitination by Slx5-Slx8 may regulate a function of Yra1 distinct from mRNA export.

### Loss of the Yra1 C-box sensitizes the genome to DSBs

Since our data indicate that Yra1 is modified by Slx5-Slx8, a STUbL important for genome stability (39), we examined whether the abrogation of Yra1 ubiquitination induces defects in genome integrity. For this purpose, we used the *HA-yra1allKR* mutant that cannot be ubiquitinated since all the Lysines (K) are replaced by Arginines (R) **(Fig 2A)** (8).

**Fig 2:**
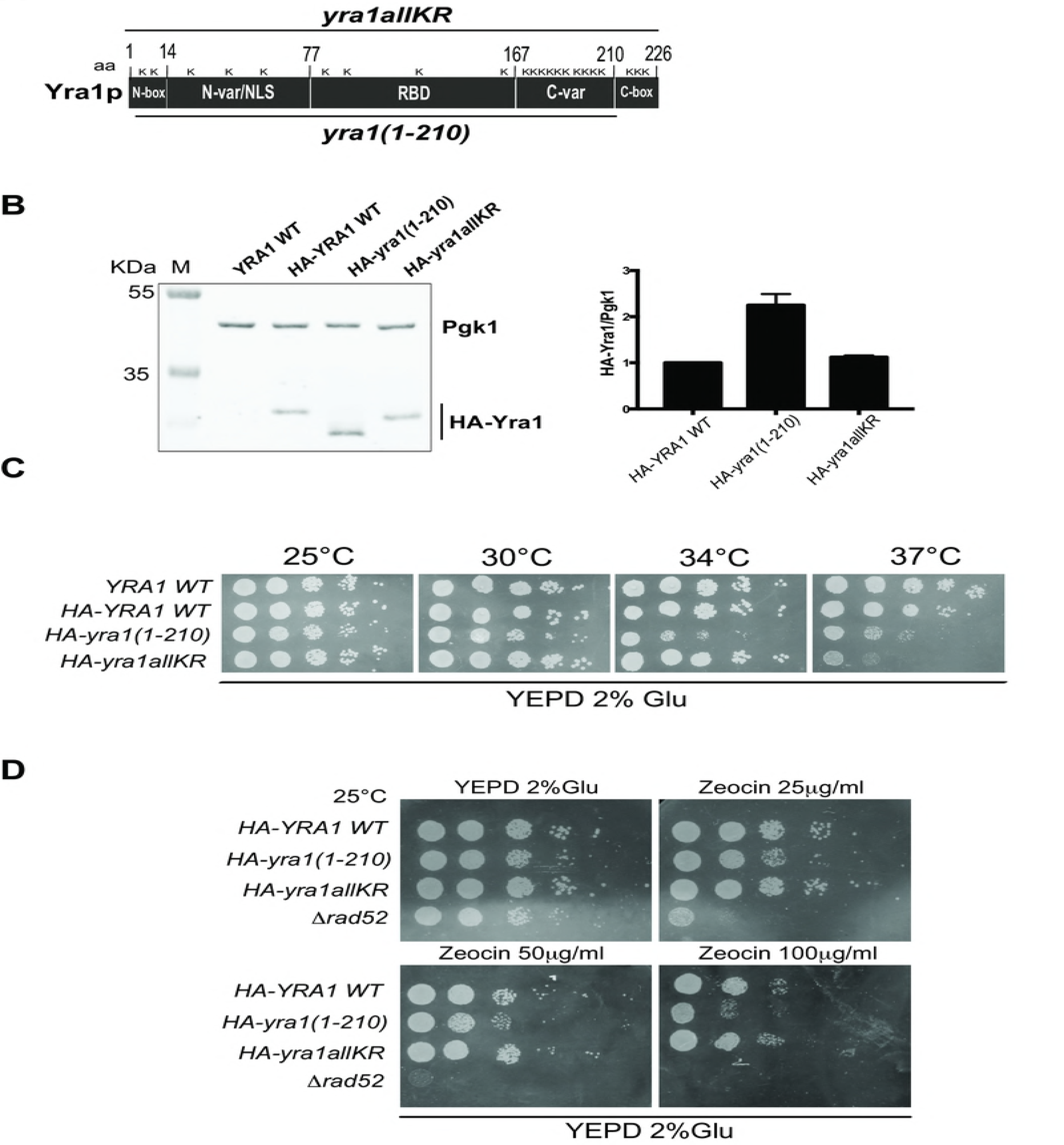
The Yra1 C-box, but not Yra1 ubiquitination, is important for genome stability. **(A)** Scheme of Yra1 mutants used in this study. **(B)** Left: Western Blot analysis of HA-Yra1 levels in integrated *HA-YRA1 WT*, *HA-yra1(1-210)*, *HA-yra1allKR*, was performed using an αHA antibody; an αPgk1 antibody was used as loading control. One representative Western blot is shown. Right: Western blot quantification showing the HA-Yra1/Pgk1 ratio of three experiments with relative standard error of the mean. The quantifications were performed using Lycor Software. **(C)** Spot test analysis of confluent cells at 25°C, 30°C, 34°C, 37°C of *YRA1 WT* (No Tag), integrated *HA-YRA1 WT*, *HA-yra1(1-210)*, and *HA-yra1allKR* strains on YEPD 2% Glucose. **(D)** Spot test analysis on YEPD 2% Glu, Zeocin 25 μg/ml, 50 μg/ml and 100 μg/ml at 25°C of confluent cells of integrated *HA-YRA1 WT* and *HA-yra1* mutants as well as *Δrad52* strains.

We also used the *HA-yra1(1-210)* mutant which codes for a protein that is still ubiquitinated (data not shown) but lacks the highly conserved 16 C-terminal amino-acids **(Fig 2A)**. Because Yra1 levels are maintained through splicing autoregulation, the intron was retained in both wild-type and mutant HA-YRA1 constructs to limit the potential toxic effect of Yra1 overexpression (4, 10, 12, 13, 16). Although the C-terminal domain has been implicated in splicing inhibition (4), the HA-yra1(1-210) protein is only mildly overexpressed compared to wild-type HA-Yra1 or the HA-yra1allKR mutant protein and presents only a slight growth defect at 25°C (**Fig 2B and C**). Both mutants are thermosensitive as shown by spot test analysis at different temperatures (25°C-30°C-34°C-37°C) (**Fig 2C**). FISH analysis did not reveal any nuclear poly(A)+ RNA retention at 25°C indicating that the two mutants have no mRNA export defect in the conditions used in this study (**Supplementary Fig S3B**) (7). Interestingly, additional spot test analyses in the presence of Zeocin, indicated that the *HA-yra1(1-210)* but not the *HA-yra1allKR* mutant is sensitive to this genotoxic drug (**Fig 2D**). This observations indicates that the Yra1 C-box is important for genome stability in the presence of DNA double strand breaks (DSBs) while Yra1 ubiquitination is not.

### Yra1 is recruited to an irreparable DSB (HO cut)

To obtain more direct evidence for a possible role of Yra1 in the DNA damage response pathway (DDR), we induced an irreparable DSB at the MAT locus using a galactose-inducible HO endonuclease as described (25) (**Supplementary Fig S4A**). Consistent with the irreparable nature of the induced HO cuts, these strains do not grow on galactose (**Supplementary Fig S5A**). Importantly, Yra1 recruitment at the HO cut, examined by ChIP with an αYra1 antibody, was significant at regions close to the DSB after 2h of HO induction (**Fig 3A**).

**Fig 3:**
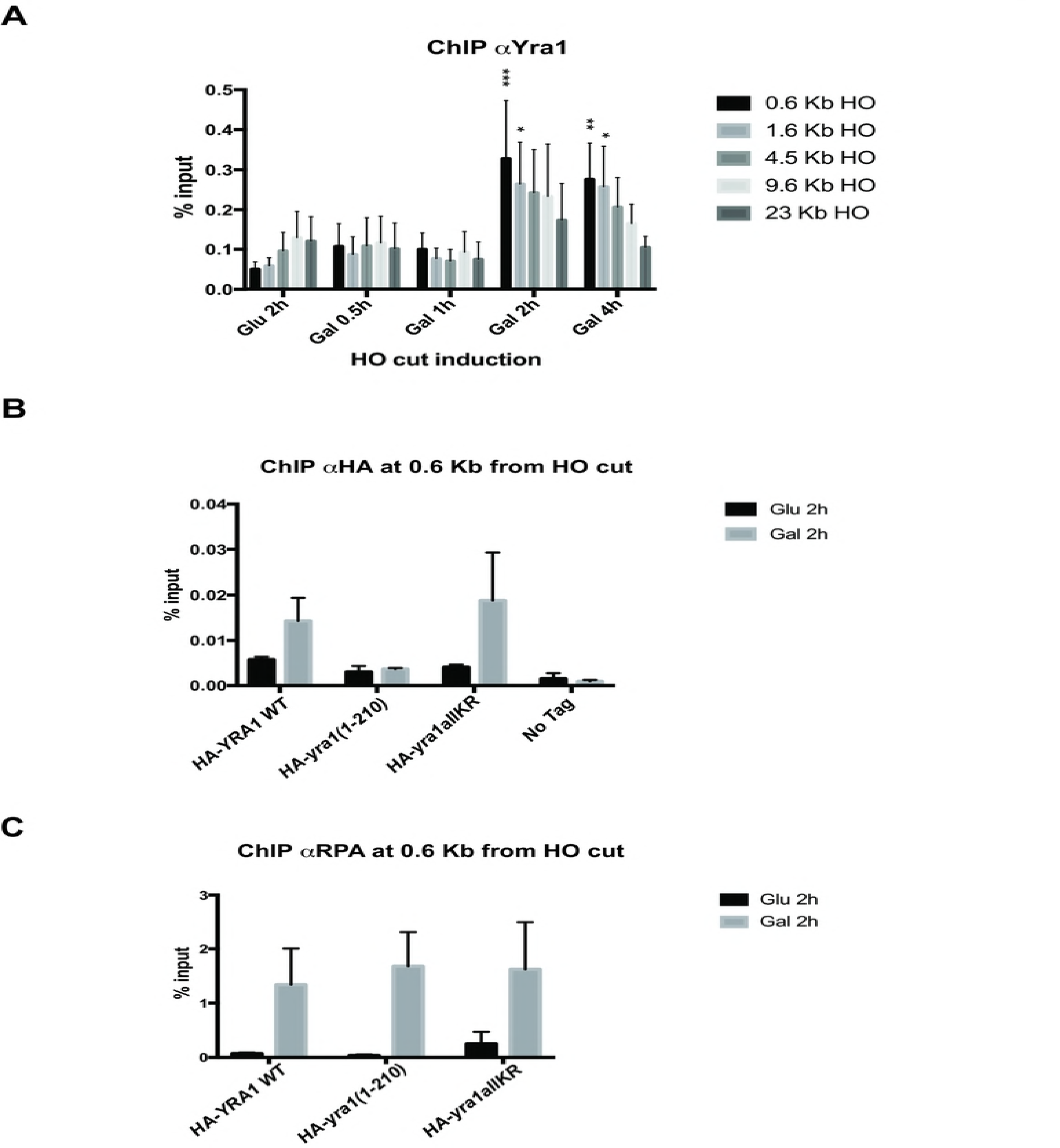
Yra1 is recruited to an irreparable DSB HO cut site. **(A)** Yra1 recruitment at the HO cut site was defined by ChIP with an αYra1 antibody after 0.5h, 1h, 2h and 4h of HO endonuclease induction with galactose using the GA6844 strain described in (25). The 2h Glucose time point was used as no cut control. ChIP values are indicated as percentage of input at 0.6Kb, 1.6Kb, 4.5Kb, 9.6Kb and 23Kb from the HO cut. The average of 6 experiments is shown with corresponding standard error of the mean. Two way ANOVA test was performed with multiple comparisons; P values < 0.05 (*), < 0.01 (**), < 0.001 (***) that refer to Glu 2h (no cut) are shown. **(B)** Yra1 mutants are differentially recruited to the irreparable HO cut site. ChIP using αHA antibody of HA-Yra1 WT, HA-yra1(1-210), HA-yra1allKR, Yra1WT (no tag) at 0.6 Kb from the HO cut site after 2h of HO induction with Galactose using the strains with *HA-YRA1 WT* or mutants integrated in strain GA6844 described in (25). The 2h Glucose time point was taken as no cut control. ChIP values are shown as percentage of input. The average of 3 independent experiments is shown with corresponding standard error of the mean. **(C)** RPA recruitment to the HO cut site in *HA-YRA1 WT* and *HA-yra1* mutants. ChIP using αRPA antibody at 0.6 Kb from the HO cut site after 2h of HO induction with Galactose in the same strains as in (B). The 2h Glucose time point was taken as no cut control. ChIP values are indicated as percentage of input. The average of 3 independent experiments is shown with corresponding standard error of the mean.

Considering that the efficiency of the cut is nearly 100% after 30’ of HO induction (25) (**Supplementary Fig S6A**), the recruitment after 2h suggests it may depend on extensive resection.

To define whether the sensitivity to Zeocin of the *HA-yra1(1-210)* mutant may be due to its impaired recruitment to DSB loci, sequences encoding HA-tagged wild-type or mutant Yra1 (*HA-YRA1 WT*, *HA-yra1(1-210)* and *HA-yra1allKR*) were integrated into the irreparable HO DSB strain at the *YRA1* locus and the recruitment of these different HA-Yra1 proteins at the HO cut was examined by ChIP using αHA antibodies after 2h in galactose (**Fig 3B**), which induces efficient HO cleavage in both wild-type and mutant strains (**Supplementary Fig S6B)**. These experiments show that the HA-yra1allKR protein is recruited to the HO cut site to similar levels as the HA-Yra1 WT in Galactose (HO cut) (**Fig 3B**). In contrast, although in this experiment the HA-yra1(1-210) protein is expressed to slightly higher levels than HA-Yra1 WT (**Supplementary Fig S5B and S5C**), its binding to the HO site does not increase in galactose, suggesting that the Yra1 C-terminal region is important for Yra1 recruitment to the DSB. Since Yra1 is recruited to the HO cut 2h after Gal induction, once there has been extensive resection, we asked whether RPA binding to the HO cut might vary in the different *HA-yra1* mutants. RPA association was not affected in the *HA-yra1* mutants despite the lack of HA-yra1(1-210) recruitment (**Fig 3C**), suggesting that RPA binding is probably not dependent on Yra1 recruitment.

### The Yra1 C-terminal region is important for DSB repair (HO cut)

Since irreparable DSBs relocate to the nuclear pore in G1/S phase (26) within 2h after cut induction (25), one possibility is that Yra1 recruitment to irreparable DSB is the consequence of HO cut re-localization to pores. To exclude this possibility, we took advantage of an HO cut reparable system (**Supplementary Fig S4B**) (28), since the DSB repair occurs within the nuclear interior (40, 41). To define whether the *HA-yra1* mutants may be defective in DSB repair, the *HA-YRA1 WT*, *HA-yra1(1-210)* and *HA-yra1ΔallKR* sequences were integrated at the *YRA1* locus of the reparable HO DSB strain and the percentage of cells surviving under HO cut induction was examined.

Three serial dilutions of exponentially growing cells were plated on galactose 2% or galactose 3%-raffinose 1% to induce the HO cut, and on glucose 2% to repress HO endonuclease expression. CFUs were counted as an indication of cells able to repair the DSB in the *HA-YRA1 WT*, *HA-yra1(1-210)*, *HA-yra1allKR* strains transformed with a pGAL-HO endonuclease plasmid or Empty Vector; a *No-Tag* and a *Δrad52* strain transformed with the Empty Vector only were used as controls as these two strains contain an endogenous pGAL-HO endonuclease sequence (**Fig 4A**).

**Fig 4:**
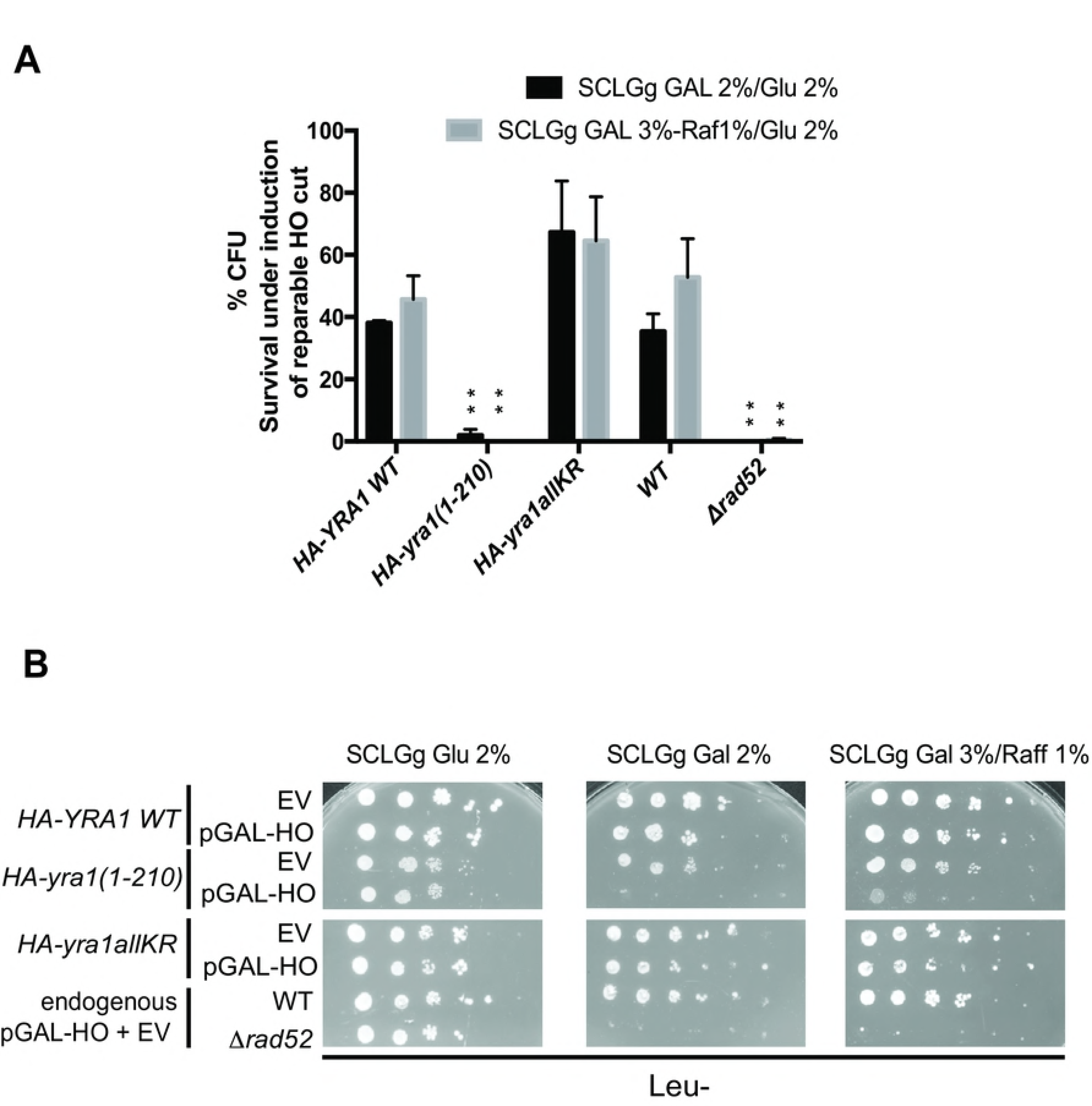
Survival under persistant induction of a reparable HO cut. **(A)** The Yra1 C-box is important for DSB repair. Three serial dilutions (200/100/50) of exponentially growing cells of NA17 strain (28) containing integrated *HA-YRA1 WT* (WT), *HA-yra1(1-210)*, *HA-yra1allKR* and transformed with pGAL-HO endonuclease containing plasmid or Empty Vector, as well as of No-Tag (NA17) and *Δrad52* strains containing endogenous pGAL-HO endonuclease and transformed with an Empty Vector were prepared. Diluted cells were plated on SCLGg Gal 2% or Gal 3%-Raf1% to constantly induce HO cut, and on SCLGg Glu 2% to repress HO endonuclease expression. The percentage of colonies was determined as the relative number of Colony Forming Units (CFUs) in each strain plated on SCLGg Gal 2% or Gal 3%-Raf1% compared to the one plated on SCLGg Glu 2%. To normalize the variability in growth due to the different media condition, the CFUs of each strain transformed with pGAL-HO endonuclease were normalized to the corresponding strain transformed with the empty vector (%CFU= (%CFU on SCLGg Gal 2% pGAL-HO/EV)/ (% CFU on SCLGg Glu 2% pGAL-HO/EV). The average of 3 independent experiments for each condition SCLGg Gal 2%/ Glu 2% and SCLGg Gal 3%-Raf 1%/ Glu 2% is shown with corresponding standard error of the mean. One way ANOVA test was performed with multiple comparisons and P value < 0.001 (**) is shown on the graph referring to *HA-YRA1 WT*. **(B)** Yra1 C-box is important for DSB repair. Spot test analysis on Leu-SCLGg Glu 2%, SCLGg Gal 2%, SCLGg Gal 3%-Raf 1%, at 25°C of exponentially growing *HA-YRA1 WT* (WT), *HA-yra1(1-210)*, *HA-yra1allKR* (transformed with pGAL-HO endonuclease containing plasmid or Empty Vector); No-Tag and *Δrad52* strains (containing endogenous pGAL-HO endonuclease sequence and transformed with an empty vector) served as controls. One representative experiment out of 3 is shown.

Interestingly, like the *Δrad52* control strain, the *HA-yra1(1-210)* mutant was not able to grow on galactose when the reparable HO cut is induced, indicating that the Yra1 C-box is important for DSB repair. This effect was confirmed by spot test analysis (**Fig 4B**). Moreover, although both *HA-yra1* mutants have comparable cut efficiency after 2h in Galactose (**Supplementary Fig S6C**), the *HA-yra1ΔallKR* showed no growth defect under HO cut induction whether in the CFU assay or the spot test, indicating that Yra1 ubiquitination is not required for DSB repair (**Fig 4A and 4B**).

Overall these observations support the view that Yra1 is important for DSB repair in a process dependent on the 16 amino acids C-terminal region. Absence of this domain may result in the inability to repair HO cuts possibly because of the reduced capacity of Yra1 to interact with the DSB.

## DISCUSSION

This study strengthens the importance of Yra1 in genome stability. In particular, our data provide evidence that the Yra1 C-terminal box is crucial for DSB repair. We have started to investigate the sensitivity of *yra1* mutants to DNA damage based on the observation that Yra1 is not only sumoylated by Siz1-Siz2 but also ubiquitinated by Slx5-Slx8, a SUMO-dependent E3 ligase important for genome stability (**Fig 1**). However, our data indicate that Yra1 ubiquitination is not important for DSB repair since the *HA-yra1allKR* mutant that completely abrogates Yra1 ubiquitination (8) does not display any genetic instability phenotypes.

To investigate the effect of Yra1 on genome stability, we rather took advantage of the *HA-yra1(1-210)* mutant that lacks the Yra1 C-box domain (**Fig 2**) without any obvious mRNA export defect (**Supplementary Fig S3**). Interestingly, our data show that the *HA-yra1(1-210)* mutant is sensitive to the DSB inducing genotoxic agent Zeocin (**Fig 2**). In line with these results, it was recently published that the DAmP allele of *YRA1* is specifically sensitive to Zeocin (18). Overall, these observations suggest that lack of the Yra1 C-box or reduced levels of Yra1 either promote DSBs or impair DSB repair.

A recent study has revealed that Npl3, an RNA binding protein involved in mRNP biogenesis, contributes to DSB resection by ensuring efficient production of *EXO1* mRNA (42). While Npl3 was proposed to have an indirect role in repair, our observations indicate that Yra1 is recruited to an irreparable DSB after 2h of cut induction and therefore extensive resection, consistent with a direct role of Yra1 in DSB repair (**Fig 3B**). Importantly, the recruitment to an irreparable DSB does not depend on Yra1 ubiquitination but requires the conserved C-box suggesting this domain may be involved in repair, although it has no effect on RPA binding to the locus (**Fig 3B and 3C**). However, we cannot fully exclude that Yra1 recruitment to irreparable DSBs may be the consequence of HO cut re-localization to the nuclear pore that occurs within 2h after cut induction (25). Furthermore, we also examined whether the irreparable DSB can be repaired by alternative pathways such us Non Homologous End Joining (NHEJ) (19) or Break Induced Repair (BIR) (26) by inducing the HO cut in the *HA-yra1* mutants for 2h and plating the cells on Glucose. The *HA-yra1allKR* and *HA-yra1(1-210)* mutants showed survival rates comparable to *HA-YRA1 WT* indicating that the Yra1 C-box and Yra1 ubiquitination do not contribute to alternative repair pathways (data not shown).

To directly address DSB repair efficiency in the *HA-yra1allKR* and *HA-yra1(1-210)* mutants, we used the HO reparable system described in (28). Unfortunately, we were unable to observe significant recruitment of Yra1 to this type of DSB by ChIP, probably because the HO reparable system is more dynamic (data not shown). However, an independent recent study identified Yra1 at an HO-induced reparable DSB using ChAP-MS (Chromatin Affinity Purification with mass spectrometry) (18). These data indicate that Yra1 is recruited to the DSB locus also when the HO cut is located within the nucleus (40, 43). Thus, the observed Yra1 binding at the irreparable HO cut (**Fig 3A and 3B**) may be specific rather than the indirect consequence of DSB relocalization to the nuclear periphery.

Besides detecting Yra1 at reparable DSBs, the recent study by Wang et al. (18) also shows that a Yra1 DAmP hypomorph mutant has a defect in global DSB repair following Zeocin treatment comparable to that observed in the absence of the central Rad52 repair protein. As discussed by the authors, this global effect probably results from the reduced expression of Rad51 due to defective mRNA biogenesis and export activity in the presence of low levels of Yra1. The same study investigated the importance of Yra1 in the repair of a single HO cut using the Yra1 anchor away system. These experiments were unable to demonstrate a role for Yra1 in this process possibly because the conditions used to deplete Yra1 by anchor away were not optimal.

Since irreparable DSBs lead to cell death (19), we addressed the critical role of Yra1 in DSB repair by defining the repair efficiency of the *HA-yra1allKR* and *□□□yra1(1-210)* mutants based on survival under induction of a reparable HO cut (**Fig 4**). Interestingly, while the *HA-yra1allKR* mutant has not effect, the *HA-yra1(1-210)* mutant exhibits very poor survival, comparable to that observed in *Δrad52* (**Fig 4**). Since the *HA-yra1(1-210)* strain has no obvious mRNA export phenotype and exhibits normal Rad51 levels (**Supplementary Fig S3** and **S7**), the data support the hypothesis that Yra1 may play a direct role in DSB repair and that the C-box is required for its recruitment to the damaged site (**Fig 3B**). In conclusion, one view is that C-box-dependent Yra1 recruitment is important for repair possibly by favoring optimal Rad52 action and homologous recombination at the DSB.

While our data show that Yra1 ubiquitination is not required for DSB repair, we cannot exclude that Yra1 sumoylation and ubiquitination by Slx5-Slx8 may facilitate relocalization of irreparable DSBs to nuclear pores (25, 26). The physiological relevance of irreparable DSB relocation to the nuclear periphery is still not fully clear. It has been speculated that it leads to proteasomal degradation of DSB-bound proteins targeted by the STUbL Slx5-Slx8 (25) to induce alternative repair pathways such us Break Induced Replication (26). In that respect, our data show that ubiquitination by Slx5-Slx8 does not lead to Yra1 degradation (**Supplementary Fig S2**). Furthermore, Yra1 ubiquitination is not required for survival after irreparable DSB induction suggesting that it is not important for non-canonical repair (data not shown).

In summary, this work indicates that at physiological expression levels, Yra1 is beneficial for genome stability by facilitating the repair of DSBs in a C-box-dependent and sumoylation/ubiquitination independent manner. Future studies should address how Yra1 recruitment to DSBs may contribute to repair through homologous recombination.

## ACKNOWLEDGMENTS

We would like to thank D. Picard and T. Halazonetis as well as members of the lab for discussions; we also are grateful to S. Gasser and M. Kupiec for strains and plasmids.

## AUTHOR CONTRIBUTIONS

V.I., E.T., and BP designed and performed experiments. N.Y.M, A.Z, V.G.M and G.S. performed experiments. V.I. and F.S. conceived experiments and wrote the paper.

---

**S1 Fig:**
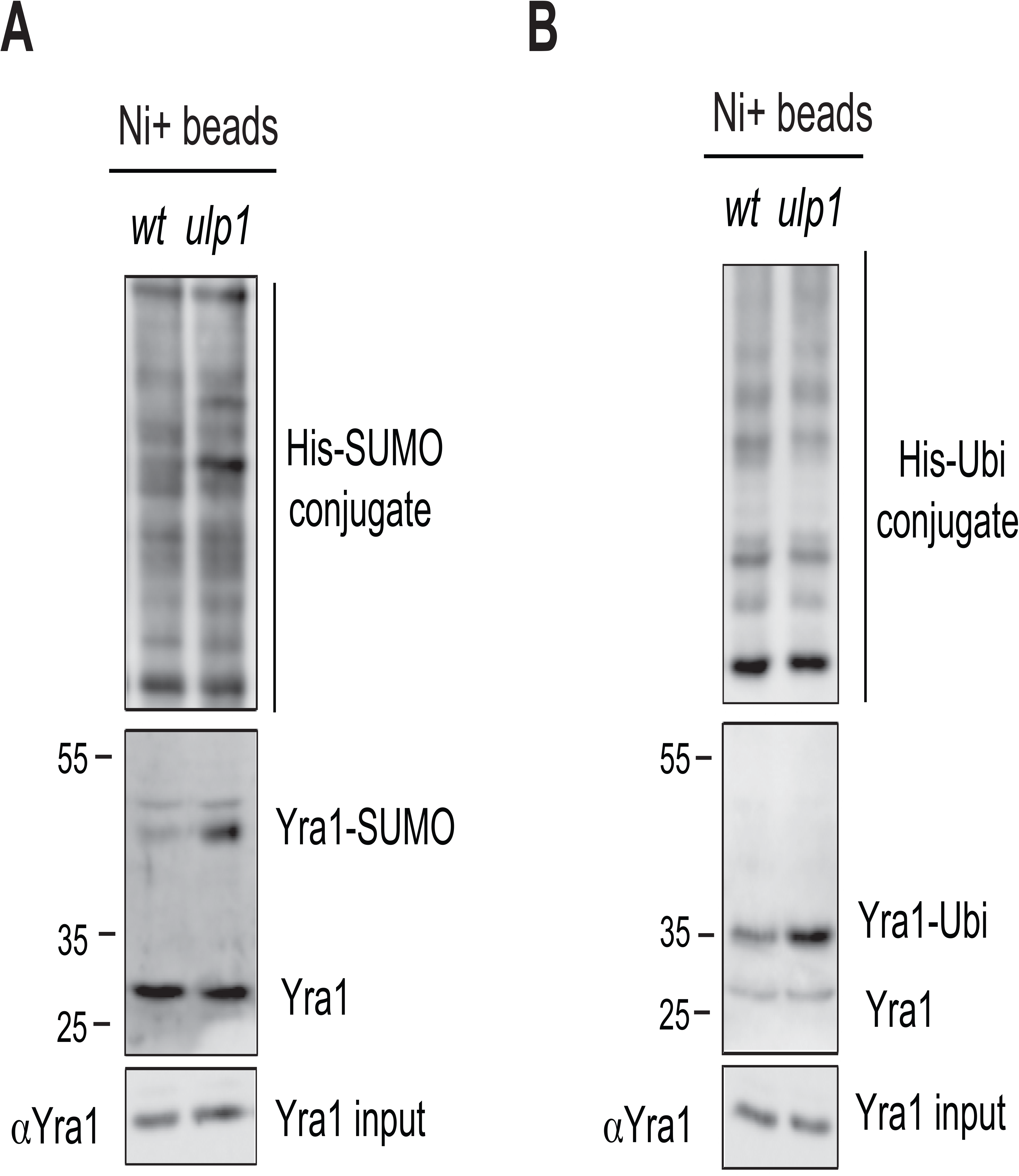
Yra1 sumoylation promotes its ubiquitination. **(A)** Yra1 is de-SUMOylated by Ulp1. Sumoylation assay in wild-type and *ulp1* temperature-sensitive (ts) mutant as described in Fig 1. One representative experiment of 3 experiments is shown. **(B)** Yra1 ubiquitination increases in the *ulp1 ts* mutant. Ubiquitination assay in wild-type and *ulp1* temperature-sensitive mutant as described in Fig 1. One representative experiment of 3 experiments is shown.

**S2 Fig:**
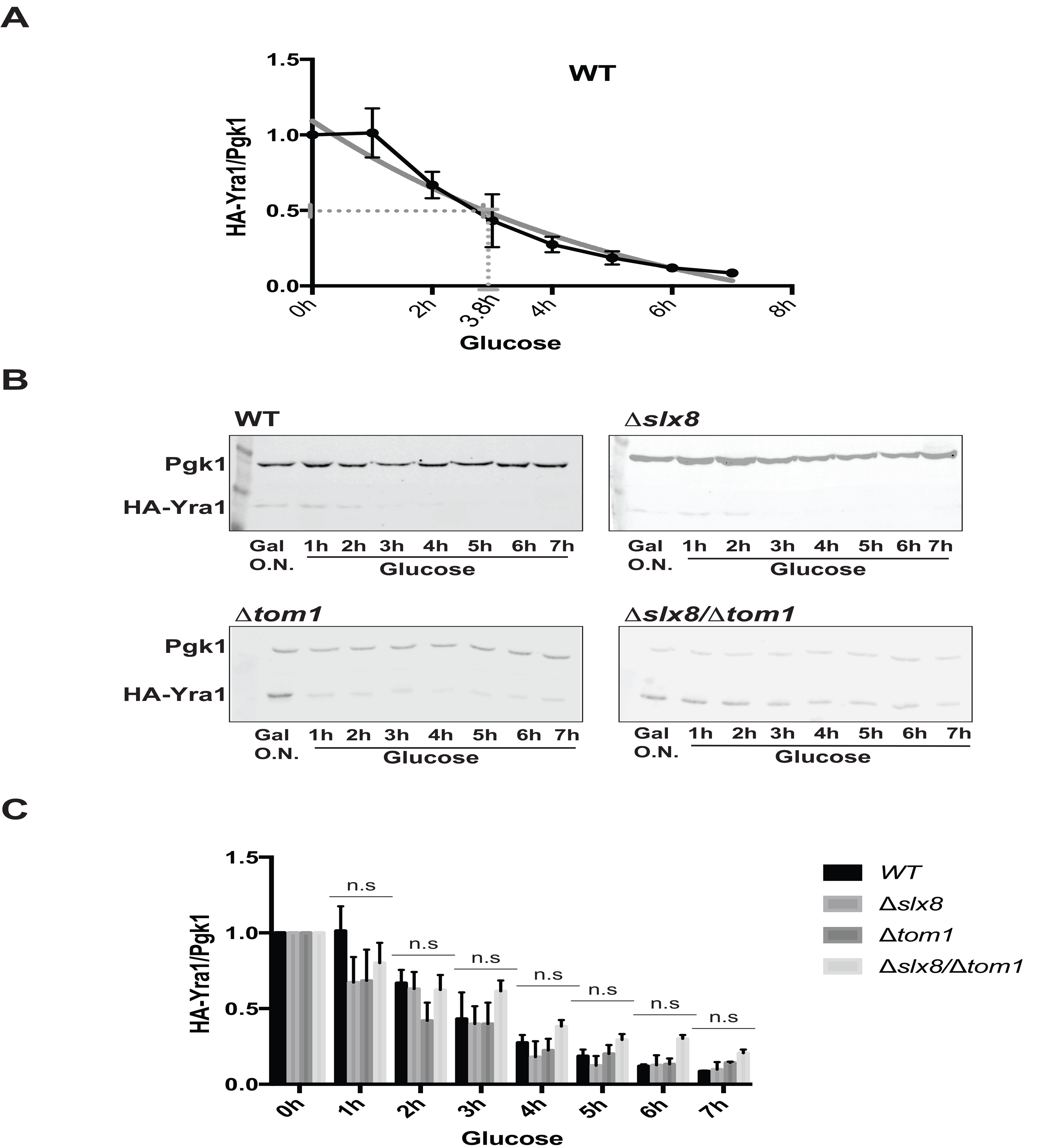
Ubiquitination by Slx5-Slx8 does not affect Yra1 half-life. **(A)** Yra1 half-life is 3.8 h when using a metabolic Gal depletion assay. Protein stability assay using metabolic depletion of *GAL-HA-YRA1* in the presence of the endogenous wild-type *YRA1* gene. *YRA1* WT shuffle cells expressing HA-Yra1 from the GAL promoter on a centromeric plasmid were grown over-night in selective medium containing 2% galactose. Cells at OD=0.3 were shifted to selective medium containing 2% glucose to repress *GAL-HA-YRA1* and collected at time 0, 1h, 2h, 3h, 4h, 5h, 6h, and 7h following glucose addition. HA-Yra1 protein levels were quantified by Western blot with an αHA antibody and normalized to Pgk1 with αPgk1 as loading control, using fluorescent secondary antibodies detected with the Lycor machine and analyzed with LITE software. The average of 2 independent experiments is shown. **(B)**, **(C)** Yra1 stability does not change in the absence of E3 ligases. Protein stability assay using metabolic depletion of *GAL-HA-YRA1* in the *YRA1* WT shuffle background combined with *Δslx5*, *Δslx8*, *Δtom1* and *Δslx8Δtom1*. Western Blot analysis (B) and relative quantification (C) were performed as described in (A). The average of 3 independent experiments (N3) is shown for the *Δslx5*, *Δslx8*, *Δtom1* and *Δslx8Δtom*1 strains. Two way ANOVA statistical test with multiple comparisons did not show any statistically significant difference (n.s) between different yeast strains in the same time points.

**S3 Fig:**
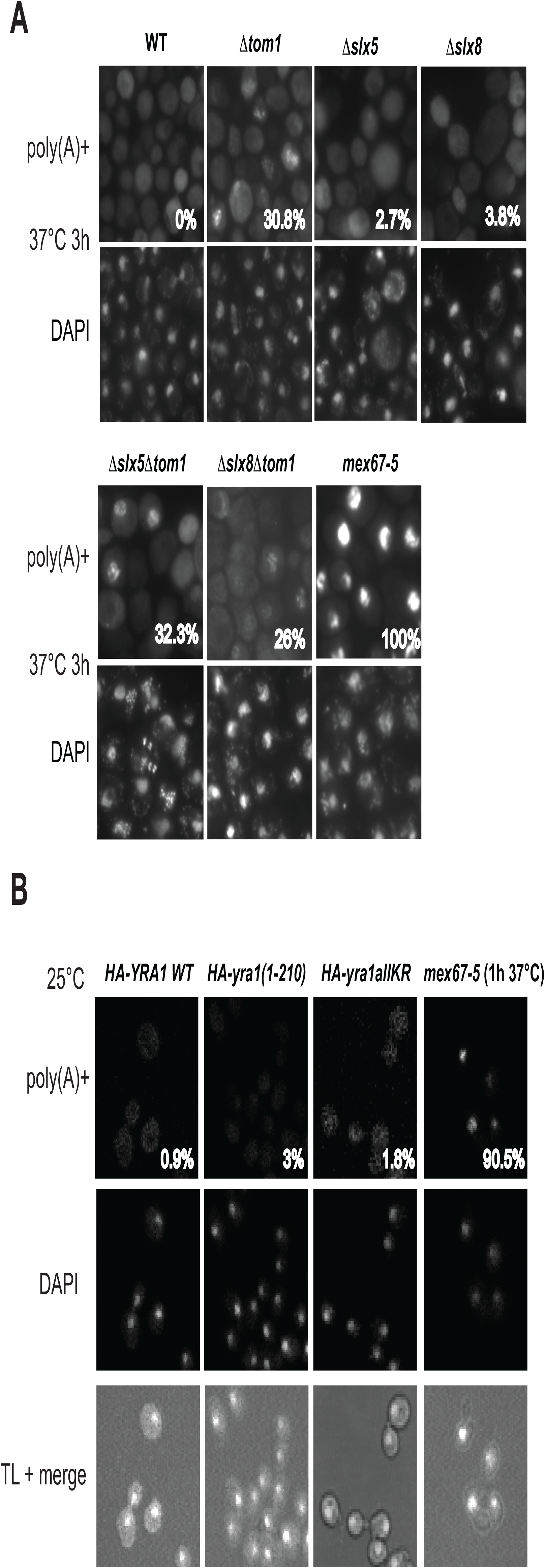
The *Δslx5*, *Δslx8* and *yra1* mutants show no mRNA export defect by FISH analysis. **(A)** Fluorescent in situ hybridization (FISH) analysis of poly(A)^+^ RNA localization using oligo(dT) probes in shuffled *HA-YRA1 WT* in WT, *Δtom1*, *Δslx8*, *Δslx5*, *Δslx8Δtom1*, *Δslx5Δtom1* background and *mex67-5* cells as control for mRNA export defect. The percent of cells showing poly(A)^+^ RNA accumulation in the nucleus is indicated in each panel. DAPI stains the cell nucleus. **(B)** Fluorescent *in situ* hybridization (FISH) analysis of poly(A)^+^ RNA localization using oligo(dT) probes in integrated *HA-YRA1* WT, *HA-yra1(1-210)*, *HA-yra1allKR* and *mex67-5* cells. Cells were grown exponentially in YEPD 2% Glu at 25°C. *mex67-5* ts mutant was grown for an additional 1h at 37°C. One representative image of nuclear staining (DAPI), oligo-dT Cy3 (poly(A)+ RNA), and Transmission Light with merged channels is shown for each strain analyzed. The percent of cells showing poly(A)^+^ RNA accumulation in the nucleus is shown in each panel.

**S4 Fig:**
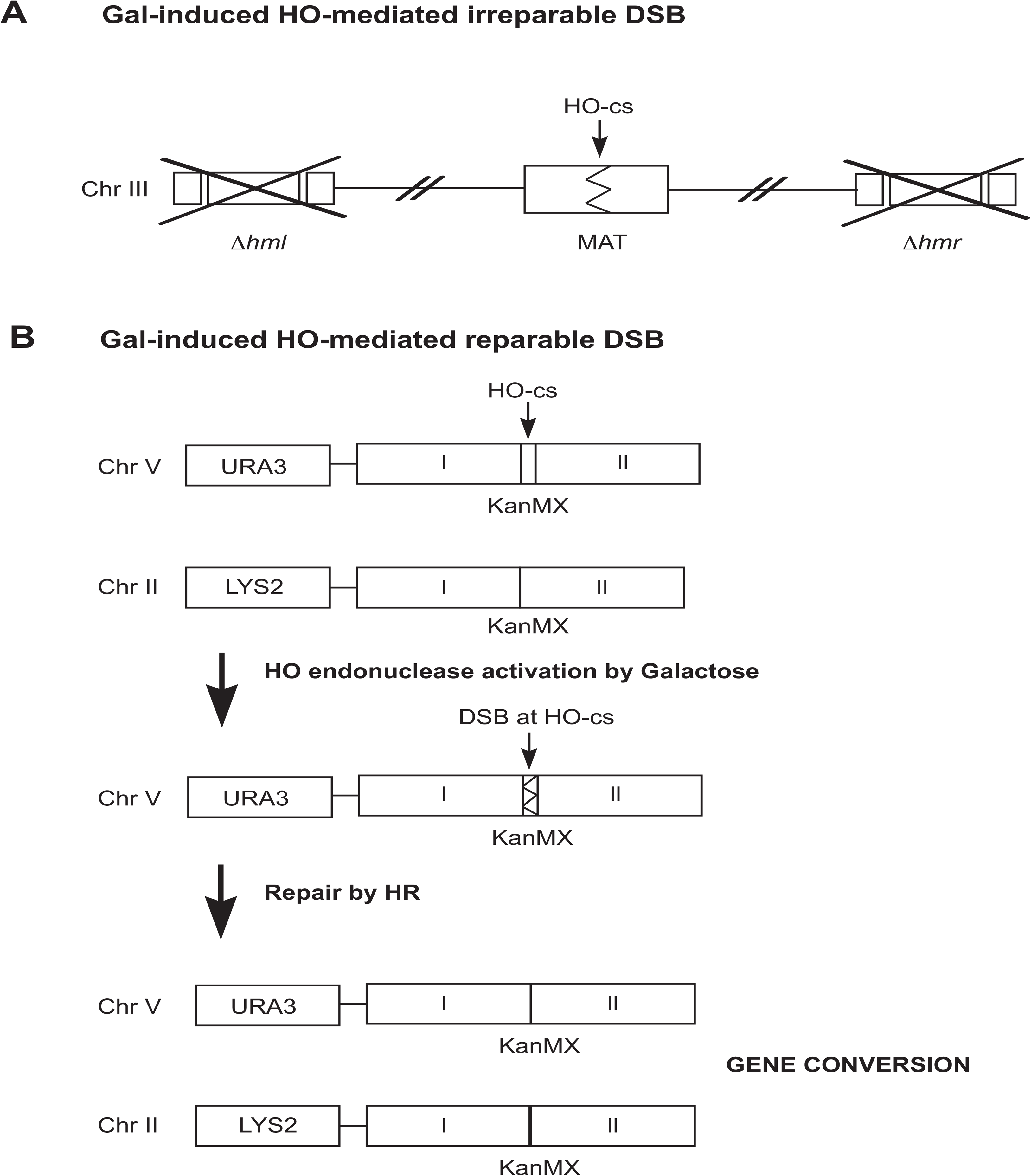
Gal-induced HO-mediated irreparable and reparable DSB systems. **(A)** Scheme showing the Gal-induced HO-mediated irreparable DSB described in (27). The HO endonuclease is expressed in the presence of Galactose, inducing the HO cut at the Mat locus that cannot be repaired because of the deletion of *HML* and *HMR*. **(B)** Scheme showing the Gal-induced HO-mediated reparable DSB described in (28). The HO endonuclease is expressed in the presence of Galactose, inducing the HO cut at the KanMx cassette next to the URA3 locus. The repair of the DSB at the HO cut is possible by HR thanks to the KanMX cassette at the LYS2 locus. If this occurs, the repair will result in an HO insensitive KanMX cassette at the URA3 locus as well as the loss of the short unique sequence surrounding the initial HO cut site.

**S5 Fig:**
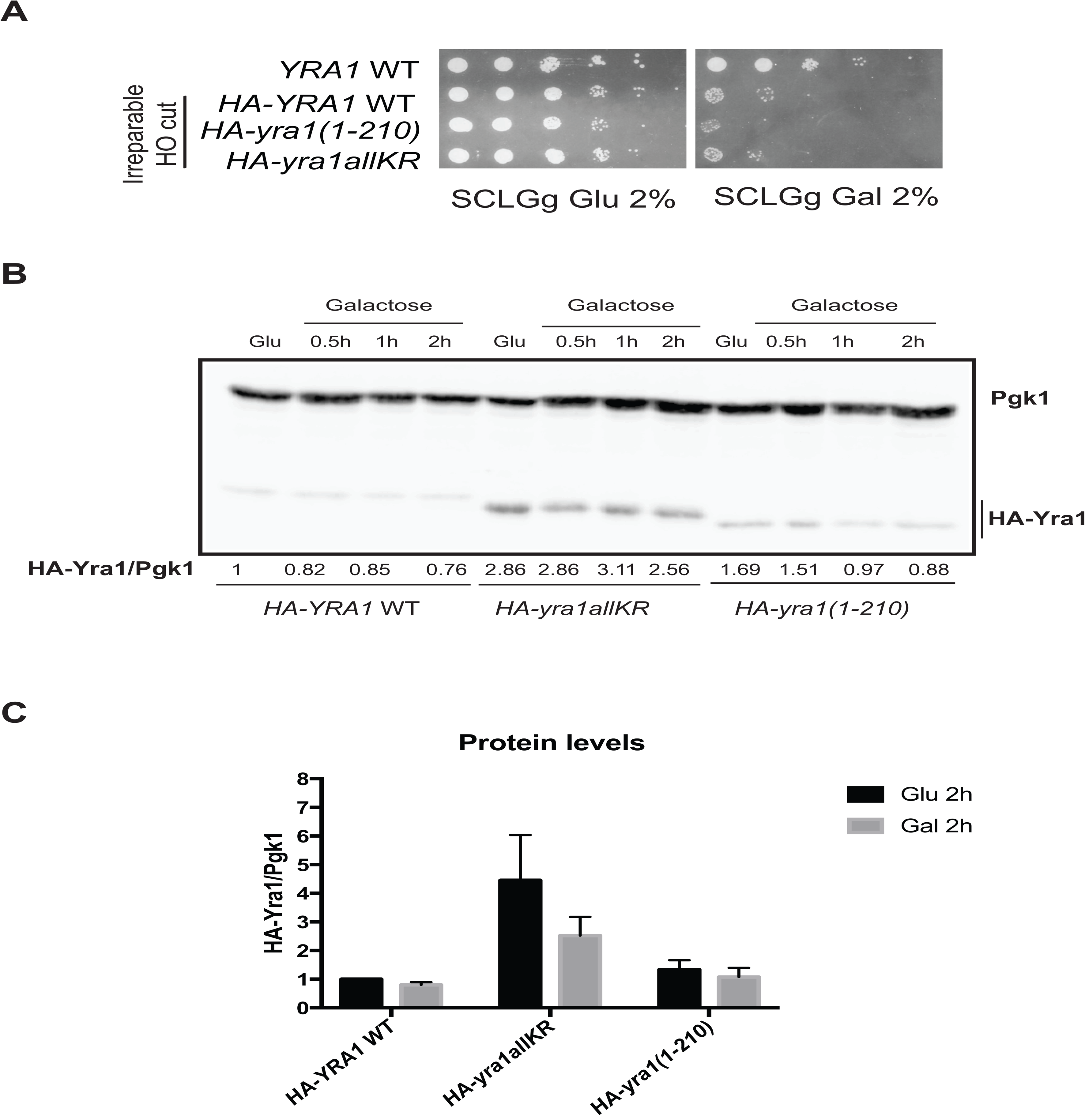
Growth phenotypes and protein levels in the irreparable HO-cut *HA-YRA1 WT* and *HA*-*yra1* mutant strains. **(A)** Spot test analysis on plates containing SCLGg Glu 2% and SCLGg Gal 2% of confluent cultures of integrated *HA-YRA1 WT* and *HA*-*yra1* mutants containing the HO irreparable DSB. A *YRA1* WT strain without any galactose-inducible irreparable HO cut is shown as control. **(B)** Protein levels of HA-Yra1 WT, HA-yra1(1-210) and HA-yra1allKR expressed from copies integrated into the GA-6844 strain (25) after 2h in Glucose or Galactose to induce the irreparable HO cut. Yra1 proteins were detected with an αHA antibody and values normalized to Pgk1 protein levels. The levels of WT or mutant HA-Yra1 proteins remain quite constant between the different time points Glu 2h, Gal (0.5h, 1h, 2h). Values of HA-Yra1/Pgk1 are shown below the blot. One representative Western Blot is shown. **(C)** Quantification of the Western blot. The average of 3 independent experiments is shown with corresponding standard error of the mean. HA-Yra1 protein levels were normalized to HA-Yra1 WT in Glu 2h set to 1.

**S6 Fig:**
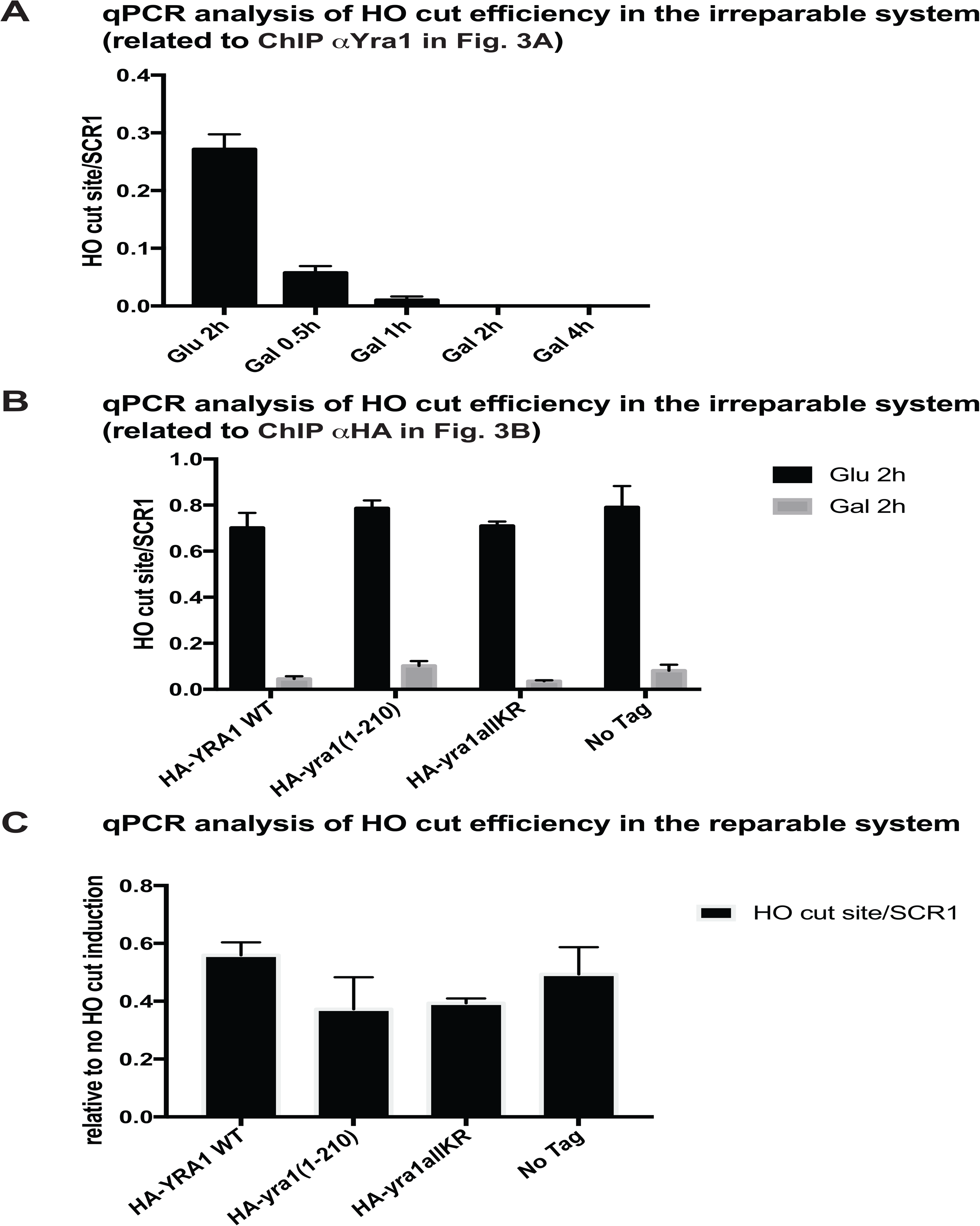
Yra1 mutants have comparable HO cut efficiency in the irreparable and repairable systems. **(A)** Analysis of HO cut site levels in the GA6844 strain described in (25) after 0.5h, 1h, 2h and 4h of HO endonuclease induction with galactose. The genomic locus was quantified by qPCR (with oligos OFS2682 + OFS2683) and the level was normalized to *SCR1*. The average of 6 independent experiments is shown with corresponding standard error of the mean. **(B)** Analysis of HO cut site levels in the *HA-YRA1 WT* and *HA-yra1* mutants integrated in GA6844 strain described in (25) after 2h of HO endonuclease induction with galactose or 2h in Glucose (no HO induction). The genomic locus was quantified by qPCR (with oligos OFS2682 + OFS2683) and the level was normalized to *SCR1*. The average of 3 independent experiments is shown with corresponding standard error of the mean. **(C)** Analysis of HO cut site levels in *HA-YRA1 WT* (WT), *HA-yra1(1-210)*, *HA*-*yra1allKR* and No-Tag strains treated with Galactose 2% (cut induction) or not (control) for 2h. The genomic locus was quantified by qPCR using oligos next to the HO site (OFS4188-OFS4179) and the level was normalized to *SCR1*. The HO cut levels in galactose are expressed relative to those in glucose. The average of 2 independent experiments is shown with corresponding standard error of the mean.

**S7 Fig:**
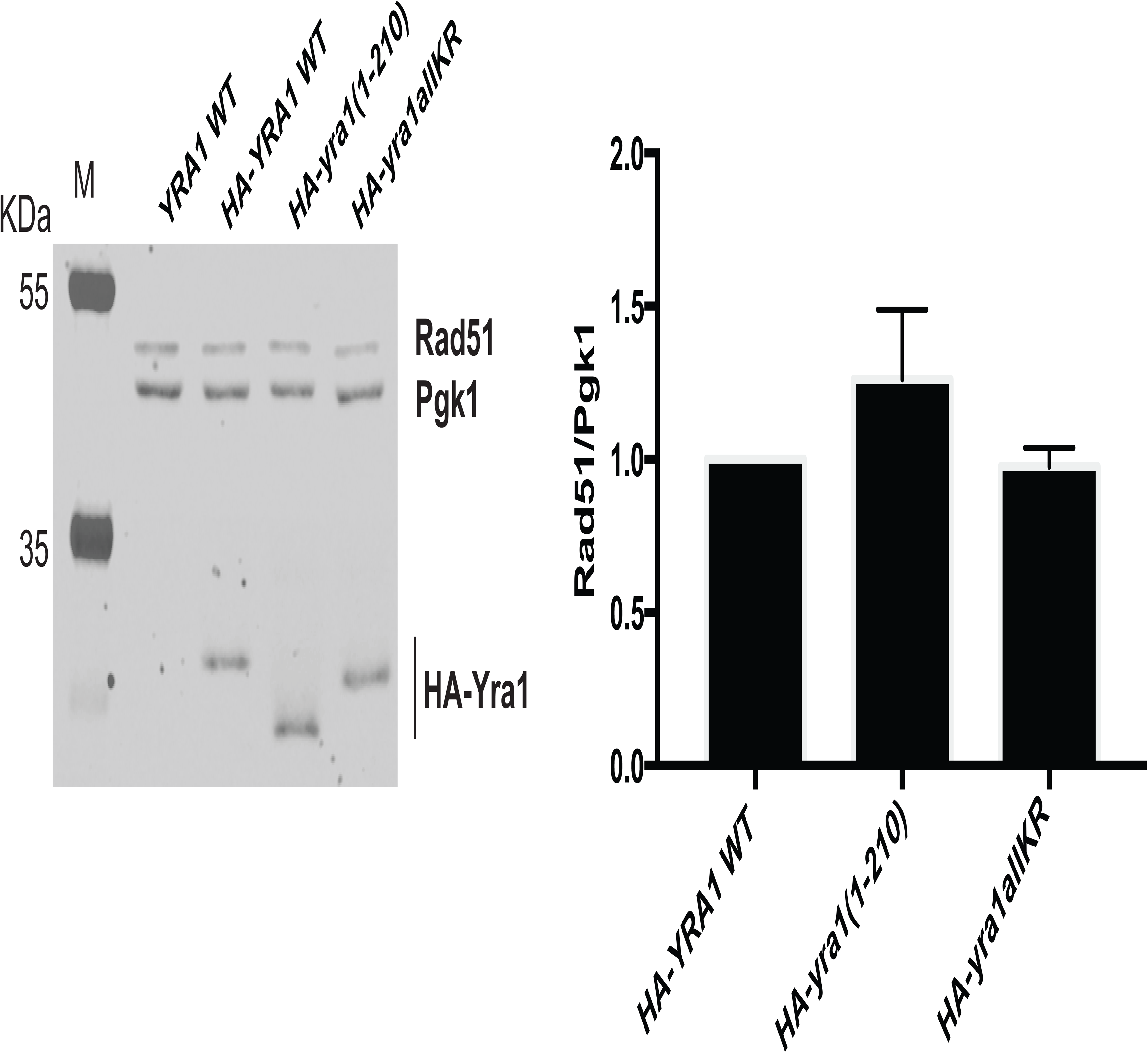
Levels of Rad51 in *HA-YRA1 WT* and *HA-yra1* mutant strains. Protein levels of Yra1 (αHA), Pgk1 (αPgk1), Rad51 (αRad51) in *HA-YRA1 WT*, *HA-yra1(1-210)* and *HA-yra1allKR* analyzed by Western blot. The right graph shows the Western blot quantification of the ratio of Rad51/Pgk1 of three independent experiments with relative standard error of the mean. The quantifications were performed using Lycor Software. Rad51 protein levels were normalized to those in *HA-YRA1 WT* that were set to 1.

**S1 Table:** Strains used in this study

| Strains | Name | Genotype | Reference |
| --- | --- | --- | --- |
| FSY1026 | *YRA1 shuffle* | *MATa ade2 leu2 trp1 ura3 Δyra1::HIS3+ <YCplac33 URA3 YRA1>* | (Zenklusen et al. 2001) |
| FSY1188 | *HA-YRA1 WT shuffled* | *MATa ade2 leu2 trp1 ura3 Δyra1::HIS3+ <YCplac22 TRP1 HA-YRA1 WT >* | This study |
| FSY4976 | *WT (W303)* | *MATa ade2 leu2 his3 trp1 ura3* | EUROSCARF |
| FSY4726 | *Δslx8 YRA1 shuffle* | *MATa ade2 leu2 trp1 ura3 Δyra1::HIS3+ <YCplac33 URA3 YRA1>, Δslx8::LEU2* | This study |
| FSY4917 | *Δslx5 YRA1 shuffle* | *MATa ade2 leu2 trp1 ura3 Δyra1::HIS3+ <YCplac33 URA3 YRA1>, Δslx5::KANr* | This study |
| FSY4934 | *Δslx5, Δslx8 YRA1 shuffle* | *MATa ade2 leu2 trp1 ura3 Δyra1::HIS3+ <YCplac33 URA3 YRA1>, Δslx5::KANr, Δslx8::LEU2* | This study |
| FSY4935 | *Δtom1, Δslx5 YRA1 shuffle* | *MATa ade2 leu2 trp1 ura3 Δyra1::HIS3+ <YCplac33 URA3 YRA1>, Δtom1::KANr, Δslx5::KANr* | This study |
| FSY4936 | *Δtom1, Δslx8 YRA1 shuffle* | *MATa ade2 leu2 trp1 ura3 Δyra1::HIS3+ <YCplac33 URA3 YRA1>, Δtom1::KANr, Δslx8::LEU2* | This study |
| FSY3373 | *Δtom1 YRA1 shuffle* | *MATa ade2 leu2 trp1 ura3 Δyra1::HIS3+ <YCplac33 URA3 YRA1>, Δtom1::KANr* | (Iglesias et al. 2010) |
| FSY3412 | *Δtom1 HA YRA1 WT shuffled* | *MATa ade2 leu2 trp1 ura3 Δyra1::HIS3+ <YCplac22 TRP1 HA-YRA1WT>, Δtom1::KANr* | This study |
| FSY4753 | *Δslx8 HA YRA1 WT shuffled* | *MATa ade2 leu2 trp1 ura3 Δyra1::HIS3+ <YCplac22 TRP1 HA-YRA1WT>, Δslx8::LEU2* | This study |
| FSY4937 | *Δslx5 HA YRA1 WT shuffled* | *MATa ade2 leu2 trp1 ura3 Δyra1::HIS3+ <YCplac22 TRP1 HA-YRA1WT>, Δslx5::KANr* | This study |
| FSY4938 | *Δslx5, Δslx8 HA YRA1 WT shuffled* | *MATa ade2 leu2 trp1 ura3 Δyra1::HIS3+ <YCplac22 TRP1 HA-YRA1WT>, Δslx5::KANr, Δslx8::LEU2* | This study |
| FSY4939 | *Δtom1, Δslx5 HA YRA1 WT shuffled* | *MATa ade2 leu2 trp1 ura3 Δyra1::HIS3+ <YCplac22 TRP1 HA-YRA1WT>, Δtom1::KANr, Δslx5::KANr* | This study |
| FSY4941 | *Δtom1, Δslx8 HA YRA1 WT shuffled* | *MATa ade2 leu2 trp1 ura3 Δyra1::HIS3+ <YCplac22 TRP1 HA-YRA1WT>, Δtom1::KANr, Δslx8::LEU2* | This study |
| FSY50410 | *Δsiz1 (YV1168)* | *MATa ade2 leu2 trp1 ura3 his3 RAD5 can1 Δsiz1::KANr* | B. Palancade |
| FSY5051 | *Δsiz2 (YV1169)* | *MATa ade2 leu2 trp1 ura3 his3 RAD5 can1 Δsiz2::KANr* | B. Palancade |
| FSY5052 | *Δsiz1, Δsiz2 (YV1077)* | *MATa ade2 leu2 trp1 ura3 his3 RAD5 can1 Δsiz1::KANr, Δsiz2::KANr* | B. Palancade |
| FSY5053 | *mms21-11 (YV1084)* | *MATa ade2 leu2 trp1 ura3 his3 RAD5 can1 mms21-11::KANr* | B. Palancade |
| FSY3992 | *ulp1 ts* | *MATa ade2 leu2 trp1 ura3 Δulp1::HIS3+ <YCplac22 TRP1 ulp1-ts>* | This study |
| FSY7017 | *HA-YRA1 WT integrated* | *MATa ade2 leu2 his3 trp1 ura3 HA-YRA1WT::HIS5* | This study |
| FSY7019 | *HA-yra1(1-210) integrated* | *MATa ade2 leu2 his3 trp1 ura3 HA-yra1(1-210)::HIS5* | This study |
| FSY7022 | *HA-yra1allKR integrated* | *MATa ade2 leu2 his3 trp1 ura3 HA-yra1Δintron::HIS5* | This study |
| FSY7158 | *Δrad52, HA-YRA1 WT integrated* | *MATa ade2 leu2 his3 trp1 HA-YRA1WT::HIS5, Δrad52::NATr* | This study |
| FSY1982 | *mex67-5* | *MATa ade2 his3 leu2 trp1 ura3 mex67-5 integrated* | (Jimeno et al. 2002) |
| FSY5073 | *GA-6844* | *JKM179, MATa, Δhml::ADE1 hmr::ADE1 ade3::GALHO ade1-100 leu2-3, 112 lys5 trp1::hisG ura3-52CFP-NUP49 GFP-LacI:Leu2 MAT::LacO repeats:TRP1* | (Horigome et al. 2014) |
| FSY6286 | *HA-YRA1 WT integrated in GA-6844* | *JKM179, MATa, Δhml::ADE1 hmr::ADE1 ade3::GALHO ade1-100 leu2-3, 112 lys5 trp1::hisG ura3-52CFP-NUP49 GFP-LacI:Leu2 MAT::LacO repeats:TRP1, HA-YRA1WT::URA3* | This study |
| FSY6287 | *HA-yra1(1-120) integrated in GA-6844* | *JKM179, MATa, Δhml::ADE1 hmr::ADE1 ade3::GALHO ade1-100 leu2-3, 112 lys5 trp1::hisG ura3-52CFP-NUP49 GFP-LacI:Leu2 MAT::LacO repeats:TRP1, HA-yra1(1-210)::URA3* | This study |
| FSY6288 | *HA-yra1allKR integrated in GA-6844* | *JKM179, MATa, Δhml::ADE1 hmr::ADE1 ade3::GALHO ade1-100 leu2-3, 112 lys5 trp1::hisG ura3-52CFP-NUP49 GFP-LacI:Leu2 MAT::LacO repeats:TRP1, HA-yra1allKR::URA3* | This study |
| FSY6881 | *NA17* | *MK225, MATa-inc, ade3::GALHO ade2-1 leu2-3, 112 his3-11, 15 trp1-1 can1-100, KanMX::HO-cs in URA3, KanMX::Clal in LYS2* | (Agmon et al. 2013) |
| **FSY7181** | *HA-YRA1 WT integrated in NA17* | *MK225, MATa-inc, ade2-1 leu2-3, 112 his3-11, 15 trp1-1 can1-100, KanMX::HO-cs in URA3, KanMX::Clal in LYS2, HA-YRA1WT::HIS5* | This study |
| **FSY7183** | *HA-yra1(1-120) integrated in NA17* | *MK225, MATa-inc, ade2-1 leu2-3, 112 his3-11, 15 trp1-1 can1-100, KanMX::HO-cs in URA3, KanMX::Clal in LYS2, HA-yra1(1-210)::HIS5* | This study |
| **FSY7186** | *HA-yra1allKR integrated in NA17* | *MK225, MATa-inc, ade2-1 leu2-3, 112 his3-11, 15 trp1-1 can1-100, KanMX::HO-cs in URA3, KanMX::Clal in LYS2, HA-yra1allKR::HIS5* | This study |
| **FSY7738** | *Δrad52 integrated in NA17* | *MK225, MATa-inc, ade3::GALHO ade2-1 leu2-3, 112 his3-11, 15 trp1-1 can1-100, KanMX::HO-cs in URA3, KanMX::Clal in LYS2, Δrad52::NATr* | This study |

**S2 Table:** Plasmids used in this study

| **Code** | **Name** | **Description** | **Reference** |
| --- | --- | --- | --- |
| **pFS 3087** | *pRS426 His6-UBI* | *Ubiquitin under CUP1 promoter (URA3, 2μ)* | *(Vitaliano-Prunier et al. 2008)* |
| **pFS 3571** | *pRS426 empty* | *Empty vector (URA3, 2μ)* |  |
| **pFS 3613** | *pRS426 His6-SMT3* | *Sumo under CUP1 promoter (URA3, 2μ)* | *B. Palancade* |
| **pFS 3688** | *YCpLac111 LEU2 GAL1p HA-YRA1 WT* | *YRA1 cloned as Sall fragment into pFS3647 (Gal1p HA-Sall-+500 YRA1)* | *This study* |
| **pFS 3687** | *YCpLac22 TRP1 GAL1p HA-YRA1 WT* | *YRA1 cloned as HindIII-SpeI from pFS3688 into pFS2233 (TRP1, CEN)* | *This study* |
| **pFS 1341** | *YCpLac111 LEU2* | *Empty vector (LEU2, CEN)* |  |
| **pFS 3932** | *pUC18 URA3 HA-YRA1 WT* | *pUC18 with SmaI fragment containing 500 bp YRA1 5' flanks, an ATG and HA-YRA1 WT followed by 500bp Yra1 3' flanks, URA3 marker and UTR-YDR381C sequence* | *This study* |
| **pFS 3934** | *pUC18 URA3 HA-yra1(1-210)* | *pUC18 with SmaI fragment containing 500 bp YRA1 5' flanks, an ATG and HA-yra1(1-210) followed by 500bp Yra1 3' flanks, URA3 marker and UTR-YDR381C sequence* | *This study* |
| **pFS 3933** | *pUC18 URA3 HA-yra1allKR* | *pUC18 with SmaI fragment containing 500 bp YRA1 5' flanks, an ATG and HA-yra1allKR followed by 500bp Yra1 3' flanks, URA3 marker and UTR-YDR381C sequence* | *This study* |
| **pFS 4131** | *pRS415 pGAL1-HO LEU2* | *HO endonuclease under GAL1 promoter* | *David Shore Lab* |
| **pFS 4082** | *pUC18 HIS5 HA-YRA1 WT* | *pUC18 with SmaI fragment containing 500 bp YRA1 5' flanks, an ATG and HA-YRA1 WT followed by 500bp Yra1 3' flanks, HIS5 marker and UTR-YDR381C sequence* | *This study* |
| **pFS 4083** | *pUC18 HIS5 HA-yra1(1-210)* | *pUC18 with SmaI fragment containing 500 bp YRA1 5' flanks, an ATG and HA-yra1(1-210) followed by 500bp Yra1 3' flanks, HIS5 marker and UTR-YDR381C sequence* | *This study* |
| **pFS 4085** | *pUC18 HIS5 HA-yra1ΔallKR* | *pUC18 with SmaI fragment containing 500 bp YRA1 5' flanks, an ATG and HA-yra1allKR followed by 500bp Yra1 3' flanks, HIS5 marker and UTR-YDR381C sequence* | *This study* |
| **pFS 3118** | *pUG27 HIS5* | *Empty vector (HIS5)* |  |
| **pFS 3574** | *pUC18* |  |  |

**S3 Table:**
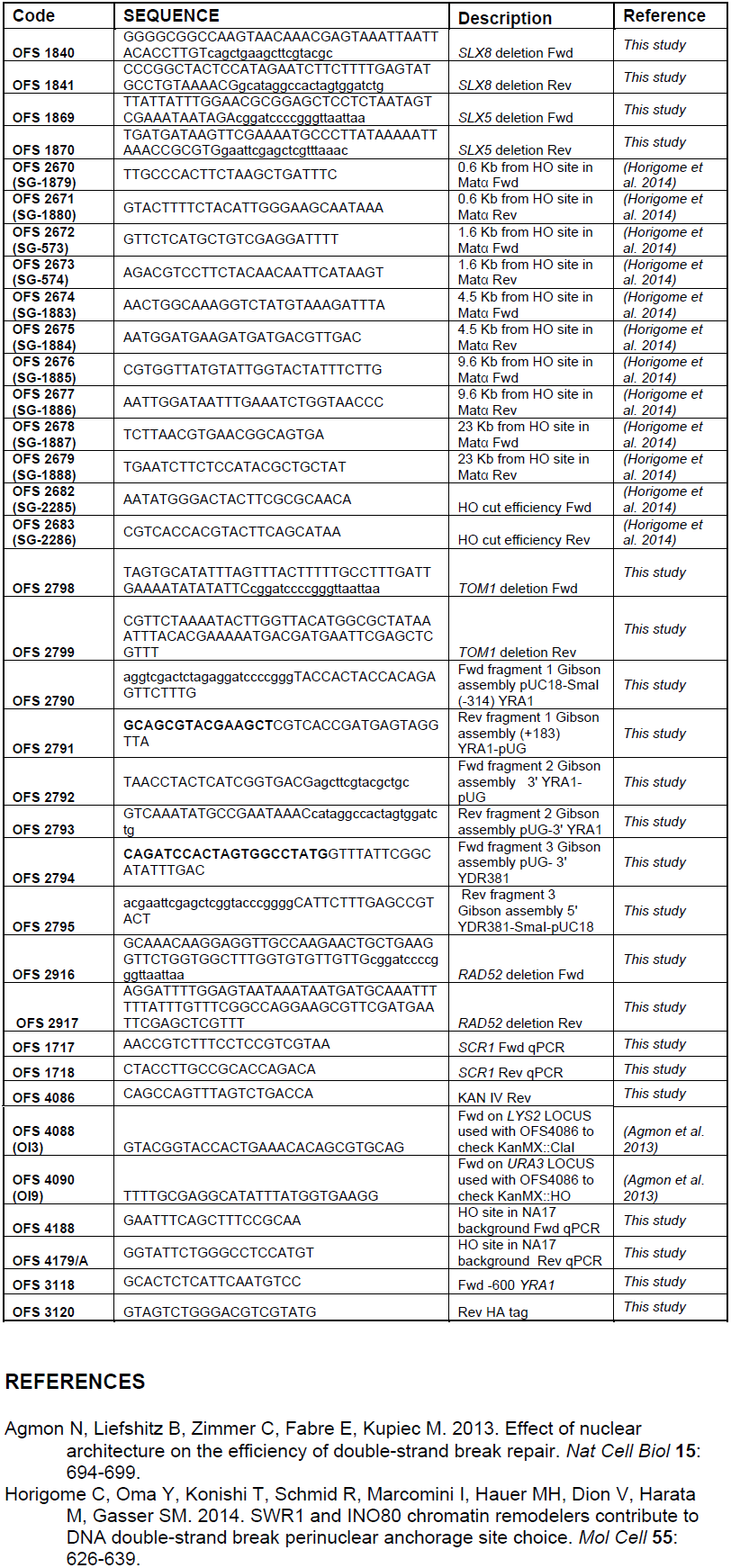
Primers used in this study

